# Comparative Analysis of In Ear and On Head EEG for Sports Applications

**DOI:** 10.64898/2026.05.07.723455

**Authors:** Ildar Rakhmatulin, Srinjoy Mitra

## Abstract

This paper presents experimental evidence that alpha-band EEG signals can be reliably detected from an in-ear electrode during physical activity, enabling fatigue monitoring in dynamic, real-world conditions such as sports. We collected an EEG dataset using a custom-designed, compact wearable system measuring only 20 mm in diameter, integrated inside the earphone. It supports five channels, four head electrodes (T3, C3, C4, T4) and one in-ear electrode, allowing simultaneous multi-site recordings. Recordings were made while a participant engaged in a controlled cycling protocol designed to induce physical fatigue. We demonstrated a direct relationship between alpha power and entropy in EEG data recorded from both the head and ear, during both activity and rest. To our knowledge, this is the first study to demonstrate in-ear alpha power tracking during active physical movement for sports-related fatigue monitoring. These findings open new possibilities for compact, wearable EEG systems in athletic and high-performance settings, where traditional EEG setups are impractical

## I. Introduction

Fatigue prediction is essential in high-stakes environments such as healthcare, transportation, and professional sports, where human performance and safety are critical. Accurate, real-time monitoring of fatigue can enable proactive interventions, reduce risk, and optimize performance. Fatigue has been linked to traffic accidents, workplace incidents, and athletic underperformance, highlighting the need for reliable detection tools.

Electroencephalography (EEG) is a promising modality for fatigue detection due to its ability to non-invasively measure brain activity with high temporal resolution. Wang et al. [1] reviewed EEG-based fatigue detection methods and emphasized their advantages in tracking physiological and cognitive states. Fatigue manifests in multiple forms: physical, mental, emotional, and chronic, which often overlap. This paper focuses on physiological fatigue, particularly in athletes, while acknowledging related influences such as stress and sleep deprivation, both of which are measurable via EEG [2– 4]. Physiological fatigue may appear in subtle forms such as reduced motivation or muscle spasms[5].

Studies have examined EEG frequency bands such as alpha, theta, beta, and gamma, to explore their correlation with performance and fatigue. For example, changes in alpha power have been linked to performance variability in elite athletes [6], while theta and Sensorimotor Rhythm (SMR) bands have been associated with attention and motor control [7]. In the context of running, EEG is used to study fatigue in real-time, especially under physical stress [8-12].

Technological advancements, particularly in machine learning, have enhanced the processing of EEG data for fatigue classification. Othmani et al. [13] highlight how machine learning methods improve EEG interpretation by extracting complex features and enabling real-time analysis. Despite this, most EEG applications remain confined to controlled environments, such as driver monitoring systems using rigid EEG caps. These setups often suffer from limitations like motion artifacts, discomfort, and a lack of practicality for long-term use.

Recent innovations in wearable EEG, notably ear-EEG devices offer more comfortable and portable alternatives. Studies have shown success in detecting drowsiness and fatigue using ear-EEG [14]. However, these devices are generally not suitable for sports activities, as they are designed for use under static conditions, and maintaining reliable contact between the electrodes and the skin becomes difficult during physical movement. Commercial interest is growing, with companies like Neurable, Emotiv, and tech giants like Apple and Meta exploring brain-monitoring technologies [15, 16].

To advance EEG-based fatigue detection, more robust systems are needed that combine wearable form factors with advanced machine learning algorithms. By leveraging signal fusion (e.g., EEG with EOG/ECG) and deep learning models such as CNNs and RNNs [17], researchers are improving the accuracy and feasibility of real-time fatigue monitoring in dynamic environments.

However, current solutions still fall short when it comes to capturing high-quality EEG data during active physical movement, particularly in sports contexts where motion artifacts and poor electrode contact undermine signal reliability. This study addresses this gap by introducing a compact, in-ear EEG system capable of detecting fatigue-related alpha-band activity during real-time exercise, offering a practical solution for mobile neuro-monitoring in athletic settings.

## II. Methos

We developed and tested a compact EEG recording device designed for fatigue detection in both resting and active states.

The prototype features a 3D-printed cap and a Brain-Computer interface. Four electrodes are placed on the scalp according to the international 10–20 system, specifically at locations T3, C3, C4, and T4, while reference and bias electrodes are integrated into the side-mounted earpieces. Both the bias and reference channels utilize four electrodes each, increasing the likelihood that at least some maintain stable skin contact, even during intense physical activity. The electrodes are made of silver/silver chloride material. The overall appearance of the prototype device is illustrated in Fig.1-a. To evaluate the potential of ear-based EEG, we added a custom electrode coated with Ag/AgCl tips inside the ear canal, Fig.1-b. This placement was tested alongside head electrodes to compare signal quality and spectral dynamics. The goal was to determine whether in-ear EEG can detect key oscillatory patterns, such as alpha rhythms.

**Fig. 1.**
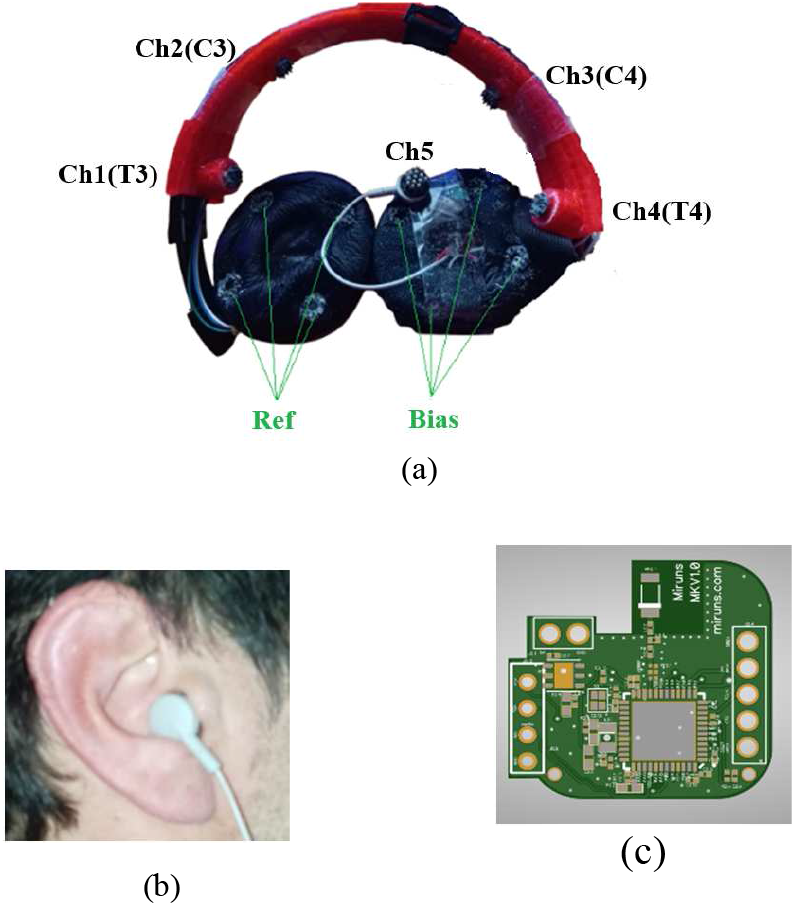
General view of the alpha rhythm setup and testing: a - the developed device with electrodes Ch1-Ch4 located on the head, Ch5 - electrode located in the ear, and reference and bias electrodes. b - ear electrode (Ch5) located in the ear, c - developed brain-computer interface, view of printed circuit board

For signal acquisition, we use an STM32WB microcontroller with Bluetooth Low Energy 5 (BLE5), enabling reliable, low-noise wireless transmission. The analog-to-digital converter (ADC) is the ADS1299 (Texas Instruments), selected for its high resolution, low noise, and integrated features suitable for EEG, such as high input impedance and bias control, Fig.1-c.

In the following, for simplicity, the channels located at T3, C3, C4, and T4 are referred to as Ch1, Ch2, Ch3, and Ch4, respectively, while the ear electrode is referred to as Ch5. We first conducted an eyes open/closed task to test alpha wave detection (∼8–13 Hz) at both the head (Ch2) and the ear (Ch5). Time-frequency analysis showed clear alpha modulation during eyes-closed intervals in both locations. While the signal amplitude was lower in the ear, the same temporal dynamics were preserved, confirming the in-ear electrode’s ability to track cognitive-state-related changes.

The alpha band comparison is visualized in Fig.2, showing strong correspondence between head and in-ear signals during the test for standard electrode location Ch2 (Fig.2-a), and for electrodes located in the ear Ch5 (Fig.2-b).

**Fig. 2.**
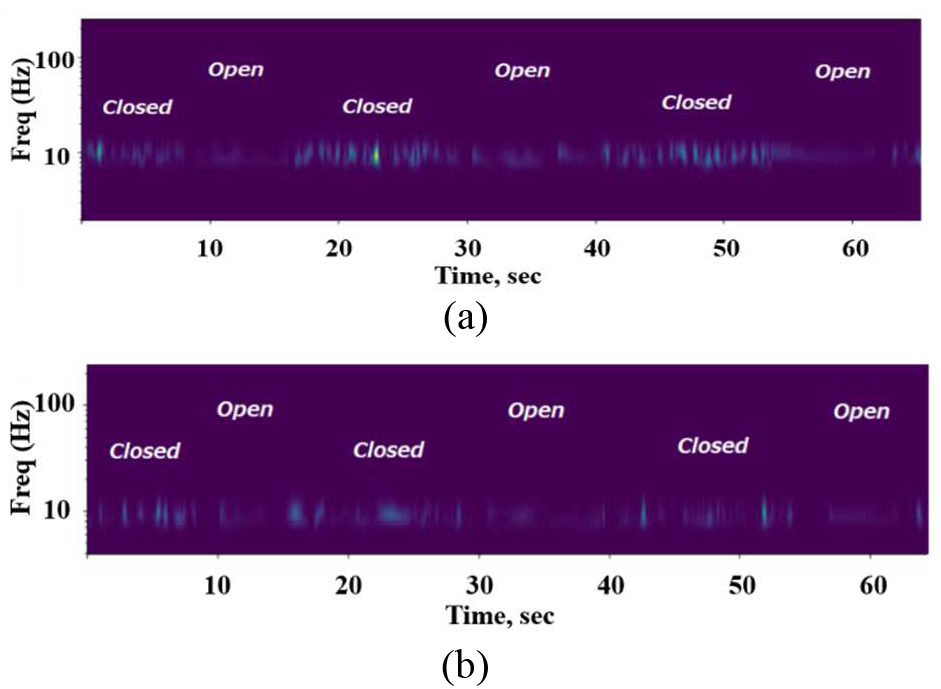
Alpha rhythm testing, a - signal in the alpha rhythm (8-12 Hz) from the electrode located on the head from Ch2, b - alpha rhythm from the ear (Ch 5) with open and closed eyes

### III. Dataset and Power in Alpha

Next, we collected EEG data from a physical activity consisting of stationary cycling. Each session included 5 minutes of pedaling and 2-minute rest intervals. This setup allowed minimal head movement and reduced motion artifacts. EEG was recorded continuously using four head electrodes (Ch1, Ch2, Ch3, Ch4) and one in-ear electrode in the left ear canal (Ch5). Electrodes were securely attached, and the participant remained seated, avoided facial movement, and did not adjust the setup during recording.

The bike’s resistance was set to a challenging level, determined through pre-testing to induce physical fatigue while maintaining data quality. Heart rate was continuously monitored and kept below 150 bpm to ensure safe aerobic intensity and avoid cardiovascular strain. Each session lasted about 40 minutes. Recordings were collected on multiple days from one healthy subject. Despite minimizing movement, some artifacts, primarily from electrode motion persisted in the data.

To clean the signals, for the collected dataset, we applied an artifact removal algorithm. Spikes exceeding 100% of average amplitude were flagged and removed across all channels. We then computed the power spectral density in the alpha band (8–13 Hz). Large spikes caused by muscle activity or motion distort EEG signals and make analysis difficult. Removing these ensures more accurate detection of alpha rhythms and cognitive states. Post-processing yielded cleaner EEG signals with clear alpha patterns, improving the detection of fatigue-related brain activity (Fig. 3).

**Fig. 3.**
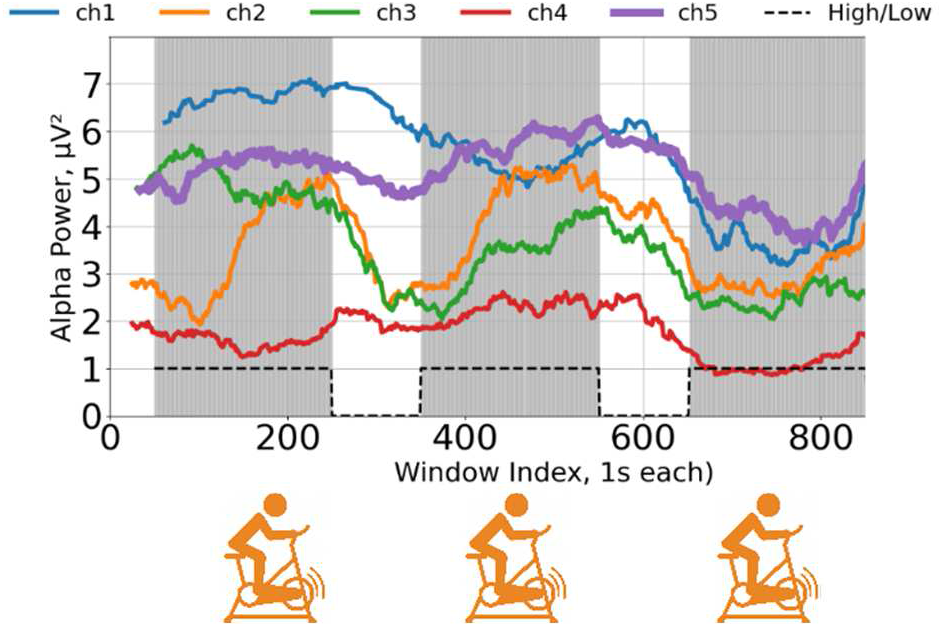
Alpha rhythm power during physical activity. Channels 1–4 represent signals from the head, and channel 5 is EEG from the ear. The black line at the bottom indicates physical activity: a high level corresponds to pedaling on the stationary bike, while a low level indicates rest

This EEG dataset records alpha power from multiple head channels (Ch1–Ch4) and one intra-ear channel (Ch5) during alternating periods of cycling and rest, marked by the black activity line. Alpha power is known to reflect cognitive and physiological states, often increasing during relaxation or fatigue and decreasing during mental or physical activity.

Across head channels, alpha power reliably increases during rest phases and decreases during active pedaling, showing a consistent and expected inverse relationship with physical effort. The synchronized behavior across these channels highlights global cortical responses to changes in physical state and suggests a robust neural signature of exertion and recovery.

Notably, the intra-ear EEG channel (Ch5) shows a similar alpha power trend reflecting rest-activity cycles, despite being located away from the head. Although the signal amplitude is slightly reduced, the alpha modulation pattern remains clear and time-locked to periods of effort and rest. This supports the feasibility of ear-EEG as a practical alternative for detecting fatigue states, especially in mobile or long-duration settings where traditional EEG setups are impractical. Taken together, these data indicate that both head and ear-EEG can track fatigue-related neural dynamics via alpha power.

## IV. Data Entropy

While traditional EEG analyses often focus on frequency-domain features—such as alpha band power—these metrics may not fully capture the nonlinear and complex neural dynamics associated with cognitive and physical fatigue. To address this limitation, we investigated the potential of entropy-based measures as complementary indicators for fatigue detection. Using the same EEG dataset that previously revealed alpha power differences between relaxed and post-exercise states, we calculated signal complexity using entropy metrics, including sample entropy and rough entropy. We calculated entropy by dividing the signal into chunks of 275 data points and measuring the uncertainty (entropy) in each chunk [18].

Our findings show a consistent decrease in entropy during physical exertion, indicating a decrease in EEG signal irregularity. This decrease suggests that fatigue may lead to more predictable or less complex neural activity, reflecting altered information processing in the brain. These observations highlight entropy as a sensitive and informative feature that complements traditional power-based approaches. By capturing temporal complexity, entropy measures provide deeper insights into neural responses to fatigue.

The results of the entropy analysis are summarized in Fig. 4, which illustrates the increase in entropy values post-exercise, aligning with the expected physiological impact of fatigue on brain dynamics.

**Fig. 4.**
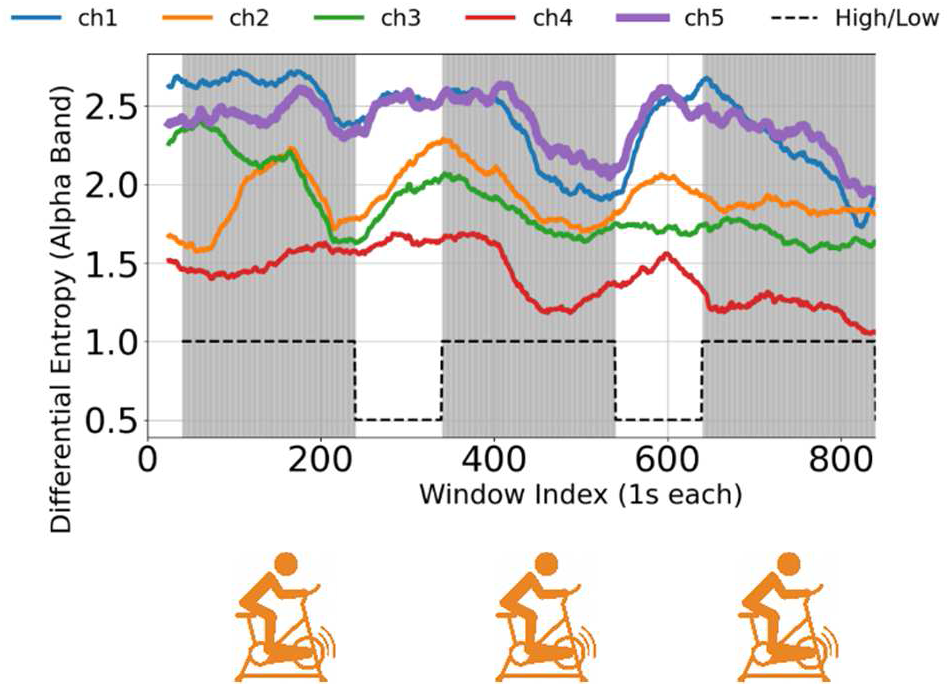
Entropy in the alpha rhythm during physical activity. Channels 1– 4 represent signals from the head, and Channel 5 is EEG from the ear. The black line at the bottom indicates physical activity: a high level corresponds to pedaling on the stationary bike, while a low level indicates rest

Entropy was calculated in fixed-size temporal windows to assess dynamic changes in signal complexity. The head electrodes showed decreased entropy during cycling, likely reflecting enhanced sensorimotor integration and neural activity under physical load. In contrast, the in-ear channel maintained a relatively high baseline entropy, which may be influenced by non-cortical factors, noise, or physiological artifacts. During rest periods, entropy values generally increased across all channels, indicating a reduction in cortical variability and complexity associated with reduced cognitive and motor demands. These findings reinforce the potential of entropy-based features as non-invasive indicators of brain state modulation during physical exertion.

The boxplot shows the distribution of alpha-band differential entropy values (8–13 Hz) across five channels (channels 1–5), with the data divided into Active and Rest states, Fig.5. The ear canal (channel 5) shows a similar pattern to the head channels: the median differential entropy is higher in the Active state than in the Rest state. This is an important finding, as it suggests that changes in the alpha band associated with changes in brain state (e.g., between Active and Rest) can also be detected using the ear EEG electrode.

**Fig. 5.**
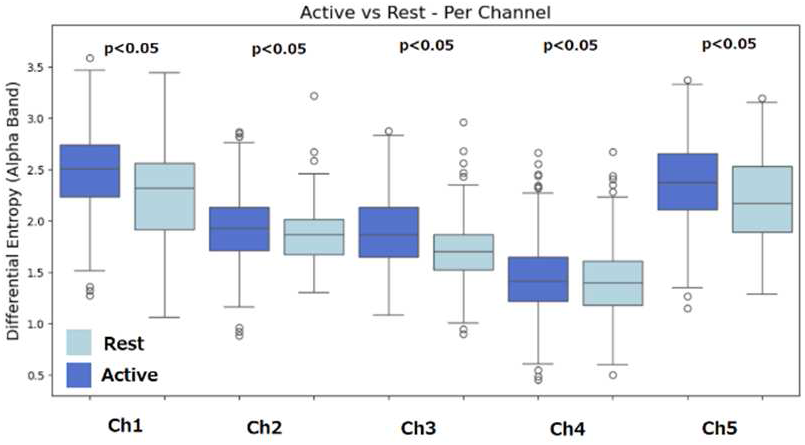
The boxplot shows the distribution of alpha-band differential entropy values (8–13 Hz) across five channels (channels 1–5)

This finding suggests that entropy reliably reflects changes in mental state across different EEG acquisition sites, including both head and in-ear recordings. Furthermore, the contrast in entropy between relaxed and fatigued states was more pronounced than the differences observed using alpha power alone. These results indicate that entropy-based analysis may serve as a more sensitive and robust marker of fatigue, particularly in setups with minimal EEG hardware, such as systems with only a few channels, and for ear EEG.

We calculated the mean (average) value for all channels, Table 1.

**TABLE I.** The mean (average) value of the EEG for rest and active conditions.

| Number of Channels | Active Mean | Rest Mean |
| --- | --- | --- |
| Ch1 | 2.47 | 2.25 |
| Ch2 | 1.92 | 1.86 |
| Ch3 | 1.9 | 1.7 |
| Ch4 | 1.42 | 1.4 |
| Ch5 | 2.37 | 2.22 |

For channels ch1, ch2, ch3, and ch5, the mean activity value is higher than the resting mean, confirming the visual trend observed in the box plot. The largest difference is observed in channel ch3, followed by ch1 and ch5. Channel ch4 shows a slight decrease in activity, which may be due to its specific location or higher sensitivity to non-brain artifacts during physical activity.

The p-value analysis shows clear differences between active and rest states in the EEG channels. Ch1, Ch2, and Ch3 display significant effects, while Ch5 exhibits a particularly strong difference, indicating that ear-EEG can reliably distinguish between active and rest conditions.

## Conclusion

This study investigated ear EEG-based fatigue detection using both alpha power and entropy as key metrics. Data were collected with a custom EEG device featuring head and in-ear electrodes. After spike artifact removal, alpha power was calculated using Welch’s method and consistently decreased after physical exertion, confirming its relevance in fatigue detection.

To enhance sensitivity, we introduced entropy analysis, which effectively captured differences between relaxed and fatigued states. Notably, similar patterns emerged in both head and ear recordings, supporting the feasibility of in-ear EEG for practical applications. Combining alpha power and entropy provided a more comprehensive and reliable approach to monitoring mental fatigue.

Ear EEG offers unique advantages for fatigue monitoring by tracking brain activity in real time. When paired with other physiological data, it enables highly personalized training, biofeedback, and recovery strategies. As wearable EEG and brain-computer interfaces evolve, they hold strong potential for broader, real-world fatigue management and cognitive performance enhancement.

This study was conducted with a single healthy participant and should therefore be considered a pilot or proof-of-concept investigation. While the results demonstrate the feasibility of monitoring exercise-induced physical fatigue using an in-ear EEG sensor during active movement, they cannot be generalized across different populations, age groups, or fitness levels. Future work will focus on expanding the study to a larger and more diverse cohort in order to establish statistical power and validate the robustness of the proposed approach.

